# Genetic architecture of host proteins interacting with SARS-CoV-2

**DOI:** 10.1101/2020.07.01.182709

**Authors:** Maik Pietzner, Eleanor Wheeler, Julia Carrasco-Zanini, Johannes Raffler, Nicola D. Kerrison, Erin Oerton, Victoria P.W. Auyeung, Jian’an Luan, Chris Finan, Juan P. Casas, Rachel Ostroff, Steve A. Williams, Gabi Kastenmüller, Markus Ralser, Eric R. Gamazon, Nicholas J. Wareham, Aroon D. Hingorani, Claudia Langenberg

**Author notes:** Correspondence to Dr Claudia Langenberg and Prof Aroon Hingorani.

## Abstract

Strategies to develop therapeutics for SARS-CoV-2 infection may be informed by experimental identification of viral-host protein interactions in cellular assays and measurement of host response proteins in COVID-19 patients. Identification of genetic variants that influence the level or activity of these proteins in the host could enable rapid ‘in silico’ assessment in human genetic studies of their causal relevance as molecular targets for new or repurposed drugs to treat COVID-19. We integrated large-scale genomic and aptamer-based plasma proteomic data from 10,708 individuals to characterize the genetic architecture of 179 host proteins reported to interact with SARS-CoV-2 proteins or to participate in the host response to COVID-19. We identified 220 host DNA sequence variants acting in *cis* (MAF 0.01-49.9%) and explaining 0.3-70.9% of the variance of 97 of these proteins, including 45 with no previously known protein quantitative trait loci (pQTL) and 38 encoding current drug targets. Systematic characterization of pQTLs across the phenome identified protein-drug-disease links, evidence that putative viral interaction partners such as MARK3 affect immune response, and establish the first link between a recently reported variant for respiratory failure of COVID-19 patients at the *ABO* locus and hypercoagulation, i.e. maladaptive host response. Our results accelerate the evaluation and prioritization of new drug development programmes and repurposing of trials to prevent, treat or reduce adverse outcomes. Rapid sharing and dynamic and detailed interrogation of results is facilitated through an interactive webserver (https://omicscience.org/apps/covidpgwas/).

## INTRODUCTION

The pandemic of the novel coronavirus SARS-CoV-2 infection, the cause of COVID-19, is causing severe global disruption and excess mortality^1,2^. Whilst ultimately strategies are required that create vaccine-derived herd immunity, in the medium term there is a need to develop new therapies or to repurpose existing drugs that are effective in treating patients with severe complications of COVID-19, and also to identify agents that might protect vulnerable individuals from becoming infected. The experimental characterization of 332 SARS-CoV-2-human protein-protein interactions and their mapping to 69 existing FDA-approved drugs, drugs in clinical trials and/or preclinical compounds^3^ points to new therapeutic strategies, some of which are currently being tested. The measurement of circulating host proteins that associate with COVID-19 severity or mortality also provides insight into potentially targetable maladaptive host responses with current interest being focused on the innate immune response^4^, coagulation^5,6^, and novel candidate proteins^7^.

Naturally-occurring sequence variation in or near a human gene encoding a drug target and affecting its expression or activity can be used to provide direct support for drug mechanisms and safety in humans. This approach is now used by major pharmaceutical companies for drug target identification and validation for a wide range of non-communicable diseases, and to guide drug repurposing^8,9^. Genetic evidence linking molecular targets to diseases relies on our understanding of the genetic architecture of drug targets. Proteins are the most common biological class of drug targets and advances in high-throughput proteomic technologies have enabled systematic analysis of the “human druggable proteome” and genetic target validation to rapidly accelerate the prioritization (or de-prioritisation) of therapeutic targets for new drug development or repurposing trials.

Identification and in-depth genetic characterization of proteins utilized by SARS-CoV-2 for entry and replication as well as those proteins involved in the maladaptive host response will help to understand the systemic consequences of COVID-19. For example, if confirmed, the reported protective effect of blood group O on COVID-19-induced respiratory failure^10^ might well be mediated by the effect of genetically reduced activity of an ubiquitously expressed glycosyltransferase on a diverse range of proteins.

In this study we integrated large-scale genomic and aptamer-based plasma proteomic data from a population-based study of 10,708 individuals to characterize the genetic architecture of 179 host proteins relevant to COVID-19. We identified genetic variants that regulate host proteins that interact with SARS-CoV-2, or which may contribute to the maladaptive host response. We deeply characterized protein quantitative trait loci (pQTLs) in close proximity to protein encoding genes, *cis*-pQTLs, and used genetic score analysis and phenome-wide scans to interrogate potential consequences for targeting those proteins by drugs. Our results enable the use of genetic variants as instruments for drug target validation in emerging genome-wide associations studies (GWAS) of SARS-CoV-2 infection and COVID-19.

## RESULTS

### Coverage of COVID-19-relevant proteins

We identified candidate proteins based on different layers of evidence to be involved in the pathology of COVID-19: 1) two human proteins related to viral entry^11^, 2) 332 human proteins shown to interact with viral proteins^3^, 3) 26 proteomic markers of disease severity^7^, and 4) 54 protein biomarkers of adverse prognosis, complications, and disease deterioration^4–6,12^ (**Fig. 1**). Of 409 proteins prioritised, 179 were detectable by an aptamer-based technology (SomaScan^©^), including 28 recognised by more than 1 aptamer (i.e. 179 proteins recognised by 190 aptamers) and 32 also measured using the Olink^©^ proximity extension assay in a subset of 485 Fenland study individuals (**Supplemental Tab. S1**). Of these 179 proteins, 111 (**Supplemental Tab. S1**) were classified as druggable proteins, including 32 by existing or developmental drugs^13^, and 22 highlighted by Gordon et al. as interacting with SARS-CoV-2 proteins^3^. To simplify the presentation of results we introduce the following terminology: we define a protein as a unique combination of UniProt entries, i.e. including single proteins and protein complexes. We further define a protein target as the gene product recognised by a specific aptamer, and, finally, an aptamer as a specific DNA-oligomer designed to bind to a specific protein target.

**Figure 1.**
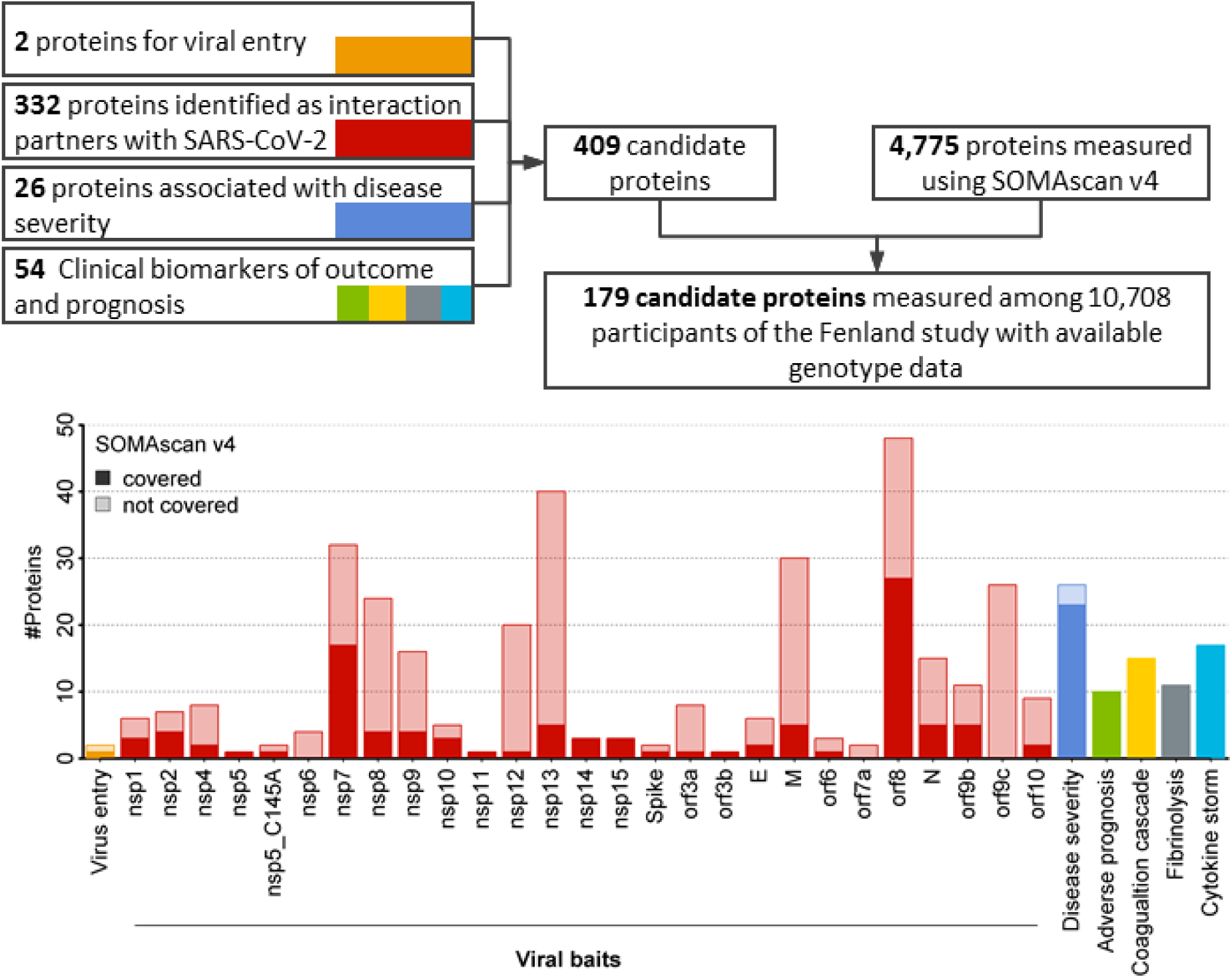
Flowchart of the identification of candidate proteins and coverage by the SomaScan v4 platform within the Fenland cohort. More details for each protein targeted are given in Supplemental Table S1.

### Local genetic architecture of protein targets

We successfully identified 220 DNA sequence variants acting in *cis* for 97 proteins recognised by 106 aptamers (**Fig. 2 and Supplemental Tab. S2**). For 45 of these proteins, no pQTLs had previously been reported. Of 9 proteins recognised by more than 1 aptamer, sentinel sequence variants were concordant (identical or in high linkage disequilibrium (LD) r^2^>0.8) between aptamer pairs or triplets for 7 proteins. Minor allele frequencies ranged from 0.01-49.9%, and the variance explained ranged from 0.3-70.1% for all *cis*-acting sentinel variants and 0.3-70.9% for *cis*-acting variants including 2-9 identified secondary signals at 57 targets, similar to what was observed considering all *cis*- and an additional 369 *trans*-acting variants identified for 98 aptamers (0.4-70.9%). Among the 97 proteins, 38 are targets of existing drugs, including 15 proteins (*PLOD2, COMT, DCTPP1, GLA, ERO1LB, SDF2, MARK3, ERLEC1, FKBP7, PTGES2, EIF4E2, MFGE8, IL17RA, COL6A1*, and *PLAT*) (8 with no known pQTL) that were previously identified^3^ as interacting with structural or non-structural proteins encoded in the SARS-CoV-2 genome and 16 proteins (*CD14, F2, F5, F8, F9, F10, FGB, IL1R1, IL2RA, IL2RB, IL6R, IL6ST, PLG, SERPINC1,SERPINE1*, and *VWF*) (7 with no known pQTL) that encode biomarkers related to COVID-19 severity^7^, prognosis, or outcome.

**Figure 2.**
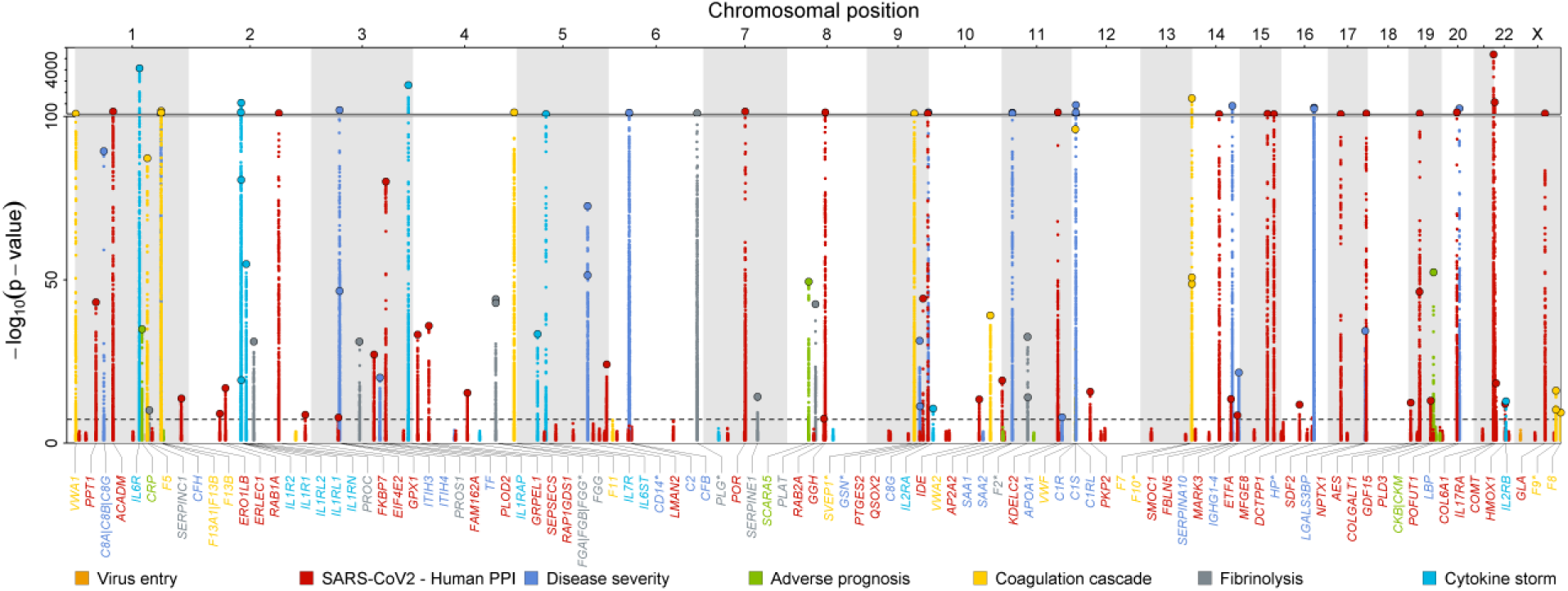
Manhattan plot of *cis*-associations statistics (encoding gene ±500kb) for 179 proteins. The most significant regional sentinel protein quantitative trait loci (pQTL) acting in *cis* are annotated by larger dots for 104 unique protein targets (dashed line; p<5×10^−8^). Starred genes indicate those targeted by multiple aptamers (n=9 genes).

Proteins are known to act in a cascade-like manner. To classify such ‘vertical’ pleiotropy, i.e. associations within a pathway, as well as ‘horizontal’ pleiotropy where proteins are acting through distinct pathways, we investigated associations of identified lead *cis*-pQTLs with all measured aptamers (N=4,776 unique protein targets, see Methods). For 38 *cis*-pQTLs mapping to druggable targets, we found evidence for a) protein specific effects for 23 regions, b) possible vertical pleiotropy for 6, and c) horizontal pleiotropy for 9 lead *cis*-pQTLs. A similar distribution across those categories was seen for the remaining *cis*-pQTLs (Fishers exact test p-value=0.49).

To test for dependencies between host proteins predicted to interact with the virus and those related to the maladaptive host response we computed genetic correlations for all proteins with at least one *cis*-pQTL and reliable heritability estimates (see **Methods**). Among 86 considered proteins, we identified a highly connected subgroup of 24 proteins including 19 SARS-CoV-2-human protein interaction partners (e.g. RAB1A, RAB2A, AP2A2, PLD3, KDEL2, GDP/GTP exchange protein, PPT1, GT251 or PKP2) and 5 proteins related to cytokine storm (IL-1Rrp2 and IL-1Ra), fibrinolysis (PAI-1), coagulation (coagulation factor X(a)), and severity of COVID-19 (GSN (gelsolin)) (**Fig. 3**). The cluster persisted in different sensitivity analyses, such as omitting highly pleiotropic genomic regions (associated with >20 aptamers) or lead *cis*-pQTLs (**Supplementary Fig. S1**). Manual curation highlighted protein modification and vesicle trafficking involving the endoplasmic reticulum as highly represented biological processes related to this cluster. Among these proteins, nine are the targets of known drugs (e.g. COMT, PGES2, PLOD2, ERO1B, XTP3B, FKBP7, or MARK3). The high genetic correlation between these proteins indicates shared polygenic architecture acting in *trans*, which is unlikely to be driven by selected pleiotropic loci identified in the present study.

**Figure 3.**
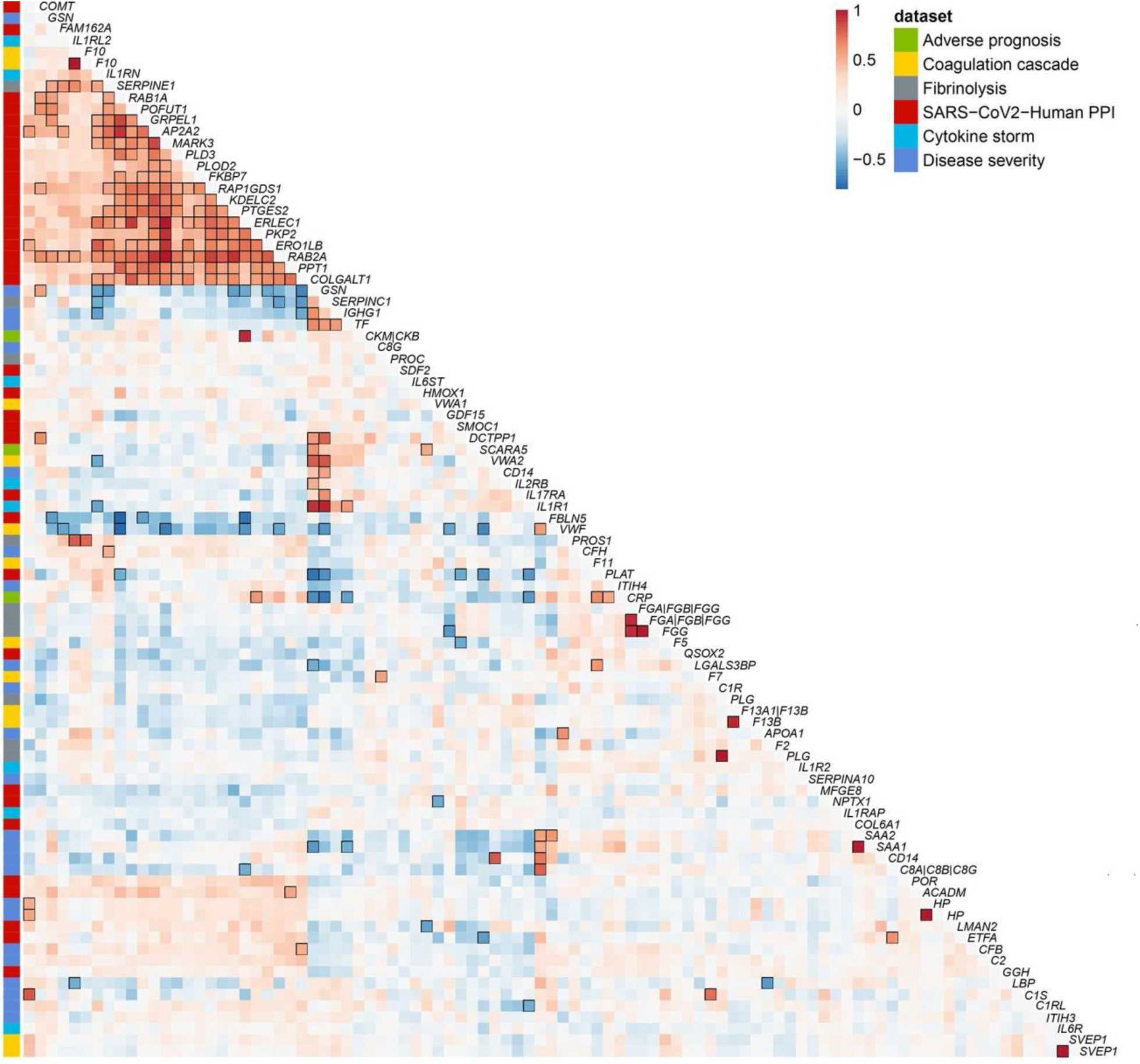
Genetic correlation matrix of 86 unique proteins targeted by 93 aptamers with reliable heritability estimates (see Methods). Aptamers were clustered based on absolute genetic correlations to take activation as well repression into account and protein encoding genes were used as labels. The column on the far left indicates relevance to SARS-CoV-2 infection. Strong correlations (|r|>0.5) are indicated by black frames.

Apart from this cluster, we identified strong genetic correlations (|r|>0.5) between smaller sets of proteins related to COVID-19 severity, and host proteins relevant to viral replication such as between IL-6 induced proteins (SAA1, SAA2, and CD14) and fibulin 5 (FBLN5).

### A tiered system for trans-pQTLs

In the absence of an accepted gold standard for the characterization of *trans*-pQTLs, we created a pragmatic, tiered system to guide selection of *trans*-pQTLs for downstream analyses. We defined as a) ‘specific’ *trans*-pQTLs those solely associated with a single protein or protein targets creating a protein complex, b) ‘vertically’ pleiotropic *trans*-pQTLs those associated only with aptamers belonging to the same common biological process (GO-term), and c) as ‘horizontally’ pleiotropic *trans*-pQTLs all remaining ones, i.e. those associated with aptamers across diverse biological processes. We used the entire set of aptamers available on the SomaScan v4 platform, N=4,979, to establish those tiers.

Among 451 SNPs acting solely as *trans*-pQTLs, 114 (25.3%) were specific for a protein target, 29 (6.4%) showed evidence of vertical pleiotropy, and 308 (68.3%) evidence of horizontal pleiotropy, indicating that *trans*-pQTLs exert their effects on the circulating proteome through diverse mechanisms. As an extreme example, the most pleiotropic *trans*-pQTL (rs4648046, minor allele frequency (MAF)=0.39) showed associations with over 2,000 aptamers and is in high LD (r^2^=0.99) with a known missense variant at *CFH* (rs1061170). This missense variant was shown, among others, to increase DNA-binding affinity of complement factor H^14^, which may introduce unspecific binding of complement factor H to a variety of aptamers, being small DNA-fragments, and may therefore interfere with the method of measurement more generally, rather than presenting a biological effect on these proteins. A similar example is the *trans*-pQTL rs71674639 (MAF=0.21) associated with 789 aptamers and in high LD (r^2^=0.99) with a missense variant in *BCHE* (rs1803274).

Sample handling is an important contributor to the identification of non-specific *trans*-pQTL associations. Blood cells secrete a wide variety of biomolecules, including proteins, following activation or release such as consequence of stress-induced apoptosis or lysis. Interindividual genetic differences in blood cell composition can hence result in genetic differences in protein profiles depending on sample handling or delays in time-to-spin. A prominent example seen in our results and reported in a previous study^15^ is variant rs1354034 in *ARHGEF3*, associated with over 1,000 aptamers (on the full SomaScan platform). *ARHGEF3* is a known locus associated with platelet counts^16^, albeit its exact function has yet to be determined, either genetically determined higher platelet counts or higher susceptibility to platelet activation may result in the secretion of proteins into plasma during sample preparation. While we report such examples, the extremely standardised and well controlled sample handling of the contemporary and large Fenland cohort has minimised the effects of delayed sample handling on proteomic assessment, as compared to historical cohorts or convenience samples such as from blood donors, evidenced by the fact that previously reported and established sample handling related loci, such as rs62143194 in *NLRP12*^15^ are not significant in our study.

Finally, for 27 out of 98 aptamers with at least one *cis*- and *trans*-pQTL, we identified no or only very weak evidence for horizontal pleiotropy, i.e. associations in *trans* for no more than 1 aptamer, suggesting that those might be used as additional instruments to genetically predict protein levels in independent cohorts for causal assessment.

### Host factors related to candidate proteins

We investigated host factors that may explain variance in the plasma abundances of aptamers targeting high-priority candidate proteins using a variance decomposition approach (see **Methods**). Genetic factors explained more variance compared to any other tested host factors for 63 out of 106 aptamers with IL-6 sRa, collagen a1(VI), or QSOX2 being the strongest genetically determined examples (**Fig. 4**). The composition of non-genetic host factors contributing most to the variance explained appeared to be protein specific (**Fig. 4**). For SMOC1 and Interleukin-1 receptor-like 1, for example, sex explained 23.8% and 17.9% of their variance, respectively, indicating different distributions in men and women. Other examples for single factors with large contributions included plasma ALT (15.4% in the variance of NADPH-P450 oxidoreductase) or age (14.2% in the variance of GDF-15/MIC-1). We observed a strong and diverse contribution from different non-genetic factors for proteins such as LG3BP, SAA, IL-1Ra, or HO-1 implicating multiple, in part modifiable, factors with independent contributions to plasma levels of those proteins.

**Figure 4.**
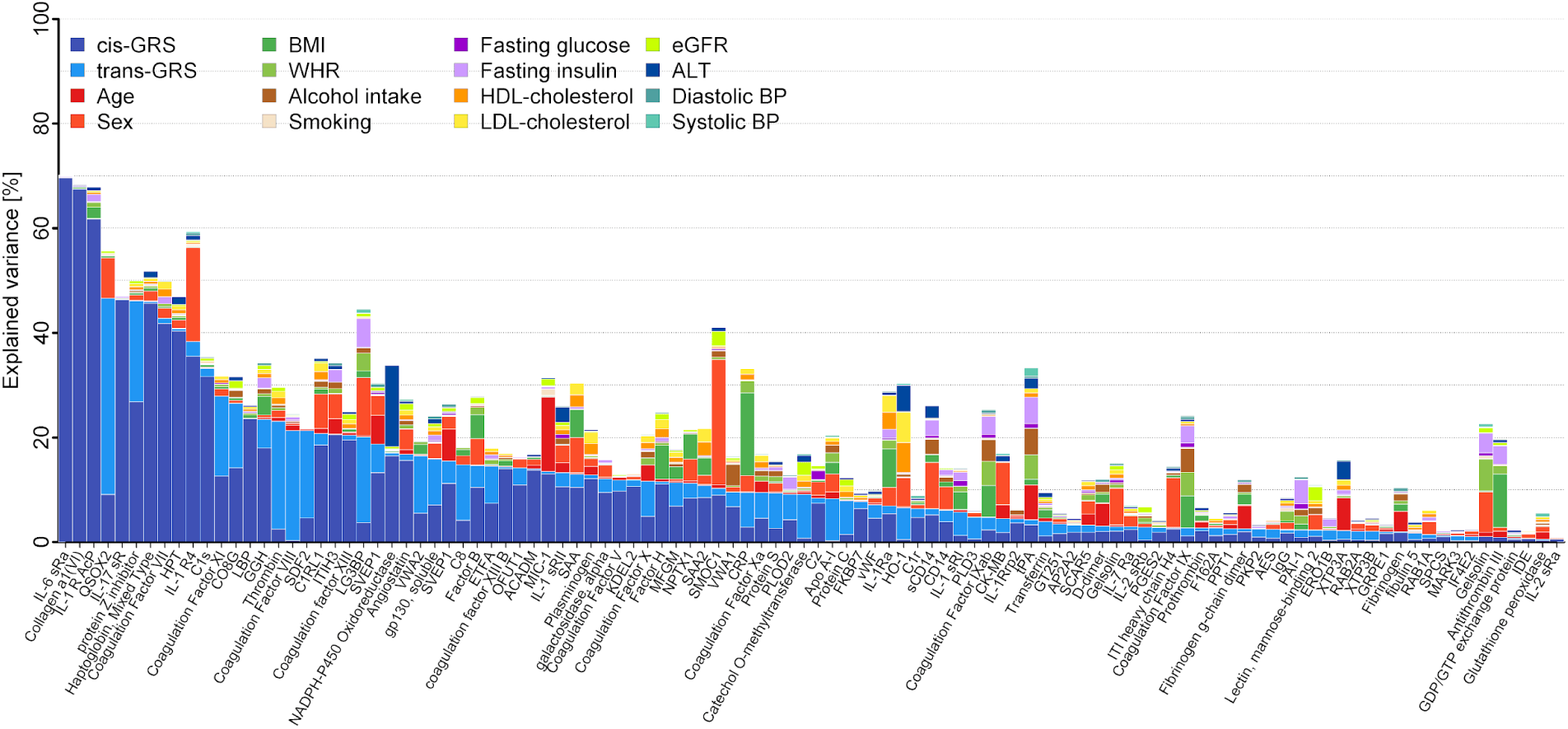
Stacked bar chart showing the results from variance decomposition of plasma abundances of 106 aptamers targeting candidate proteins. For each candidate protein a model was fitted to decompose the variance in plasma levels including all 16 factors noted in the legend. cis/trans-GRS = weighted genetic risk score based on all single nucleotide polymorphisms associated with the aptamer of interest acting in *cis* and *trans*, respectively. BMI (body mass index), WHR (waist-to-hip ratio), HDL (high-density lipoprotein), LDL (low-density lipoprotein), eGFR (estimated glomerular filtration rate), ALT (alanine amino transaminase), BP (blood pressure)

Patients with multiple chronic conditions are at higher risk of getting severe COVID-19 disease^2,17,18^ and to investigate the influence of disease susceptibility on protein targets of interest, we generated weighted genetic risk scores (GRS) for major metabolic (e.g. type 2 diabetes and body mass index (BMI)), respiratory (e.g. asthma), and cardiovascular (e.g. coronary artery disease (CAD)) phenotypes to investigate the association with all COVID-19-related proteins (**Supplemental Fig. S2**).

Plasma abundances of QSOX2 were positively associated with GRS for lung function and coronary artery disease (CAD), however, as described below these disease score to protein associations were likely driven by genetic confounding. Specifically, (*cis*) variants in proximity (±500kb) to the protein encoding gene (*QSOX2*) were genome-wide significant for forced expiratory volume (FEV1) and forced vital capacity (FVC) and exclusion of this region from the lung function genetic score abolished the score to QSOX2 association. None of the three lead *cis*-pQTLs were in strong LD with the lead lung function variant (r^2^<0.4) and genetic colocalization of QSOX2 plasma levels and lung function^19^ showed strong evidence for distinct genetic signals (posterior probability of near 100%). The association with the CAD-GRS was attributed to the large contribution of the *ABO* locus to plasma levels of QSOX2, and exclusion of this locus from the CAD score led to the loss of association with QSOX2.

The GRSs for BMI (N=10), estimated glomerular filtration rate (eGFR; N=7), and CAD (N=4) were associated with higher as well as lower abundance of different aptamers, and the asthma-GRS was specifically and positively associated with IL1RL1. Individuals with higher genetic susceptibility to BMI had higher abundances of three putative viral interaction partners (LMAN2, ETFA, and SELENOS), and lower levels of albumin, GSN, and ITIH3. Lower plasma abundances of albumin and GSN have been associated with severity of COVID-19^7^. Plasma abundance of LMAN2 (or VIP36) was associated with the BMI-GRS (positively) and the eGFR-GRS (inversely). VIP36 is shed from the plasma membrane upon inflammatory stimuli and has been shown to enhance phagocytosis by macrophages^20^. The higher plasma levels among individuals with genetically higher BMI and lower kidney function, however, do not reflect the fact that both of these are considered to be risk factors for COVID-19.

### Integration of gene expression data

We integrated gene expression data across five tissues of direct or indirect relevance to SARS-Cov-2 infection and COVID-19 (lung, whole blood, heart - left ventricle, heart - atrial appendage, and liver) from the GTEx project^21,22^ (version 8) to identify tissues and RNA expression traits contributing to protein targets. Genetically-anchored gene expression models could be established using PrediXcan^23^ for at least one of these tissues for 72 of the 102 high-priority aptamers with at least one *cis*-pQTL located on the autosomes. Protein and gene expression were significantly associated for 65 of those aptamers (p<0.05) with varying tissue specificity (Fig. 5), similar to previous reports^15,24^. Predicted gene expression (druggable targets in bold) of *ACADM*, ***SERPINC1, EROLB1***, *POR, RAB2A, KDELC2, C1RL, AES*, ***IL17RA, FKBP7***, *and* ***EIF4E2***, for example, was consistently associated with corresponding protein levels in plasma across at least three tissues, whereas gene expression in lung only was associated with plasma levels of *SAA1, SAA2*, and *SERPINA10*.

**Figure 5.**
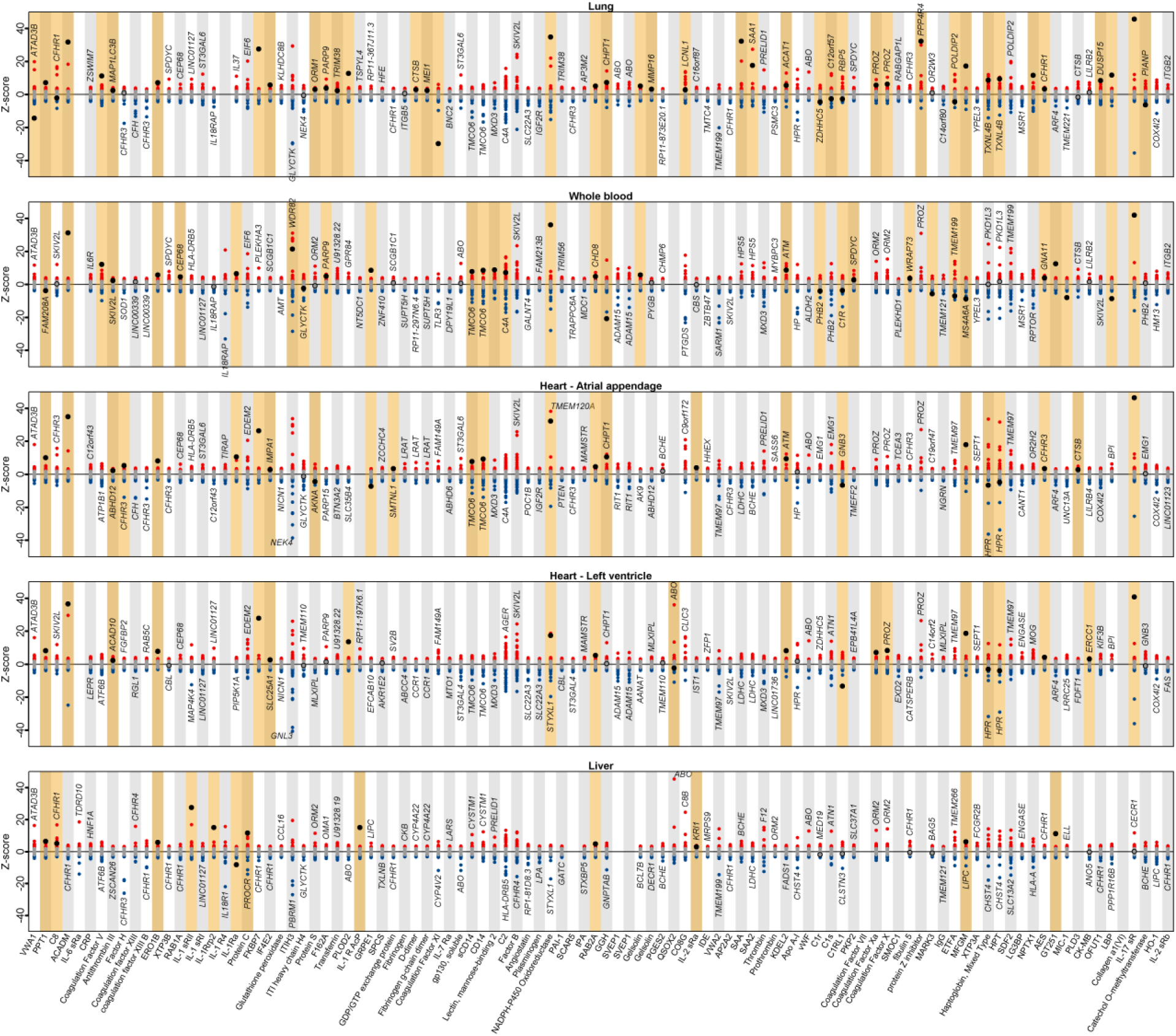
Results of predicted gene expression in each of five tissues and plasma abundances of 102 aptamers with at least one *cis*-pQTL on one of the autosomes using PrediXcan. Each panel displays results for a tissue. Each column contains results across successful gene expression models for the association with the aptamer listed on the x-axis. Red indicates nominally significant (p<0.05) positive z-scores (y-axis) and blue nominally significant inverse z-scores for associated aptamers. Protein encoding genes are highlighted by larger black circles. Orange background indicates all examples of significant associations between the protein encoding gene and protein abundance in plasma regardless if this was the most significant one. Top genes were annotated if those differed from the protein encoding gene.

Plasma levels of proteins depend on multiple biological processes rather than solely on the expression of the encoding genes. Testing for enriched biological terms^25^ across all significantly associated genes (p<10^−6^) in lung highlighted ‘signal peptide’ (false discovery rate (FDR)=2.5×10^−5^), ‘glycoproteins’ (FDR=1.7×10^−4^), or ‘disulfide bonds’ (FDR=2.8×10^−4^) as relevant processes. These are involved in the transport and posttranslational modification of proteins before secretion and highlight the complexity of plasma proteins beyond a linear dose-response relationship with tissue abundance of the corresponding mRNA.

### Cross-platform comparison

We tested cross-platform consistency of identified pQTLs using data on 33 protein targets also captured across 12 Olink protein panels and available in a subset of 485 Fenland participants. In brief, Olink’s proximity extension assays use polyclonal antibodies and protein measurements are therefore expected to be less affected by the presence of protein altering variants (PAVs) and so-called epitope effects, since they are likely to affect epitope binding only for a subset of the antibody populations, if any.

We compared effect estimates for 29 *cis*- and 96 *trans*-pQTLs based on a reciprocal look-up across both platforms (see **Methods, Supplemental Tab. S5**). We observed strong correlation of effect estimates among 29 *cis*-pQTLs (r=0.75, **Fig. S3**) and slightly lower correlation for *trans*-pQTLs (r=0.54) indicating good agreement between platforms. In detail, 36 pQTLs (30%) discovered using the far larger SOMAscan-based effort were replicated (p<0.05 and directionally consistent) in the smaller subset of participants with overlapping measurements.

We identified evidence for inconsistent lead *cis*-pQTLs for two of these 33 protein targets. The lead *cis*-pQTL for GDF-15 from SomaScan (rs75347775) was not significantly associated with GDF-15 levels measured using the Olink assay despite a clear and established signal in *cis* for the Olink measure^26^ (rs1227731, beta=0.59, p<6.5×10^−16^). However, rs1227731 was a secondary signal for the SomaScan assay (beta=0.29, p<5.8×10^−66^) highlighting the value of conditional analyses to recover true signals for cases where these are ‘overshadowed’ by potential false positive lead signals caused by epitope effects. Another protein, the poliovirus receptor (PVR), did not have a *cis*-pQTL in the SomaScan but in the Olink-based discovery (rs10419829, beta=-0.84, p<2.9×10^−33^), which in the context of an observational correlation of r=0.02 suggests that the two technologies target different protein targets or isoforms. A similar example is ACE2, the entry receptor for SARS-CoV-2, with a correlation of r=0.05 between assays and for which we identified only *trans*-pQTLs with evidence for horizontal pleiotropy (**Supplemental Tab. S3**). The SCALLOP consortium investigates genetic association data focused on Olink protein measures, and can be a useful and complementary resource for the subset of proteins of interest that are captured (https://www.olink.com/scallop/).

### Drug target analysis

We identified pQTLs for 105 proteins already the target of existing drugs or known to be druggable which are implicated in the pathogenesis of COVID-19 either through interactions with SARS-CoV-2 proteins, untargeted proteomic analysis of plasma in affected patients, or as candidate proteins in the potentially maladaptive host inflammatory and pro-coagulant responses. Of these, 18 are targets of licensed or clinical phase compounds in the ChEMBL database. Thirteen of these were targets of drugs affecting coagulation or fibrinolytic pathways and five were targets of drugs influencing the inflammatory response. Drugs mapping to targets in the coagulation system included inhibitors of factor 2 (e.g. dabigatran and bivalirudin), factor 5 (drotrecogin alfa), factor 10 (e.g. apixaban, rivaroxaban), von Willebrand factor (caplacizumab), plasminogen activator inhibitor 1 (aleplasinin), and tissue plasminogen activator. Drugs mapping to inflammation targets included toclizumab and satralizumab (targeting the interleukin 6 receptor), brodalumab (targeting the soluble interleukin-17 receptor) and anakinra (targeting interleukin-1 receptor type 1). Two targets with pQTLs (catechol O-methyltransferase and alpha-galactosidase-A) were identified as potential virus-host interacting proteins. The former is the target for a drug for Parkinson’s disease (entacapone) and the latter is deficient in Fabry’s disease, a lysosomal disorder for which migalastat (a drug that stabilises certain mutant forms of alpha-galactosidase-A) is a treatment.

Out of the 105 proteins, 24 have no current licensed medicines but are deemed to be druggable including multiple additional targets related to the inflammatory response, prioritised by untargeted proteomics analysis of COVID-19 patient plasma samples. These included multiple components of the complement cascade (e.g. Complement C2, Complement component C8, Complement component C8 gamma chain, and Complement factor H). A number of inhibitors of the complement cascade are licensed (e.g. the C5 inhibitor eculizumab) or in development, although none target the specific complement components prioritised in the current analysis.

The effect of drug action on COVID-19 for the targets identified in this analysis requires careful analysis. For example, one target identified through analysis of host-virus protein interactions is prostaglandin E synthase 2 (PGES2) involved in prostaglandin biosynthesis. Non-steroidal anti-inflammatory drugs (NSAIDs) are also known to suppress synthesis of prostaglandins and, though the evidence is weak, concerns have been raised that NSAIDs may worsen outlook in patients with COVID-19^27^. The *cis*-pQTLs we identified for PGES2 might be useful to explore this further.

### Linking cis-pQTLs to clinical outcomes

We first tested whether any of the 220 *cis*-pQTLs or proxies in high LD (r^2^>0.8) have been reported in the GWAS catalogue and identified links between genetically verified drug targets and corresponding indications for lead *cis*-pQTLs at *F2* (rs1799963 associated with venous thrombosis^28^), *IL6R* (rs2228145 with rheumatoid arthritis^29^), and *PLG* (rs4252185 associated with coronary artery disease^30^).

To systematically evaluate whether higher plasma levels of candidate proteins are associated with disease risk, we tested genetic risk scores (*cis*-GRS) for all 106 aptamers for their associations with 633 ICD-10 coded outcomes in UK Biobank. We identified 9 significant associations (false discovery rate <10%), including the druggable example of a thrombin-*cis*-GRS (2 cis-pQTLs as instruments) and increased risk of pulmonary embolism (ICD-10 code: I26) as well as phlebitis and thrombophlebitis (ICD-10 code: I80) (**Supplemental Table S6**).

To maximise power for disease outcomes, include clinically relevant risk factors, and allow for variant-specific effects we complemented the phenome-wide strategy with a comprehensive look-up for genome-wide significant associations in the MR-Base platform^31^.

Out of the 220 variants queried, 74 showed at least one genome-wide significant association, 20 of which were *cis*-pQTLs for established drug targets. We obtained high posterior probabilities (PP>75%) for a shared genetic signals between 25 *cis*-pQTLs and at least one phenotypic trait using statistical (conditional) colocalisation (**Fig. 6 and Supplemental Tab. S7**). Among these was rs8022179, a novel *cis*-pQTL for microtubule affinity-regulating kinase 3 (MARK3), a regional lead signal for monocyte count and granulocyte percentage of myeloid white cells^16^. The variant showed associations with higher plasma levels of MARK3 and monocyte count and therefore suppression of MARK3 expression with protein kinase inhibitors such as midostaurin may affect the protein host response to the virus. The important role of monocytes and macrophages in the pathology of COVID-19 has been recognised^4^, and a range of immunomodulatory agents are currently evaluated in clinical trials, with a particular focus on the blockade of IL-6 and IL-1β. Our findings indicate that proteins utilized by the virus itself, such as MARK3, SMOC1, or IL-6 receptor, may increase the number of innate immune cells circulating in the blood and thereby contribute to a hyperinflammatory or hypercoagulable state. Stratification of large COVID-19 patient populations by *cis*-pQTL genotypes that contribute to stimulation/repression of a specific immune signalling pathway is one potential application of our results. However, such investigations would need to be large, i.e. include thousands of patients, and results need to be interpreted with caution as targeting those proteins can have effects not anticipated by the genetic analysis, which cannot mimic short term and dose-dependent ‘drug’ exposure.

**Figure 6.**
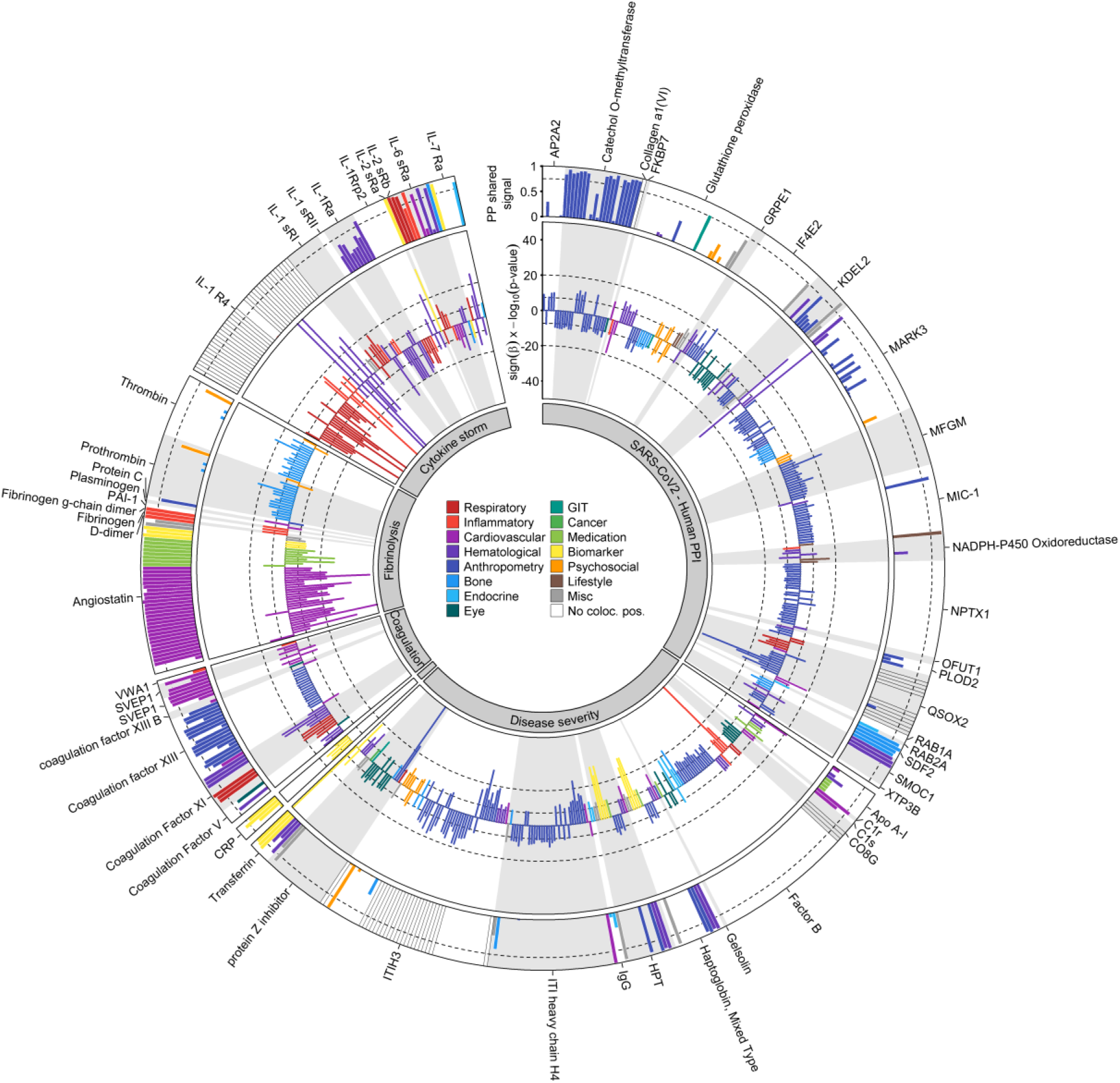
Circos plot summarizing genome-wide significant associations between 74 *cis*-pQTLs and 239 traits^31^ in the inner ring and results from statistical colocalisation in the outer ring. The dashed line in the outer ring indicates a posterior probability of 75% of shared genetic signal between the protein and a phenotypic trait. Protein targets are classified on the basis of their reported relation to SARS-CoV-2 and COVID-19. Each slice contains any *cis*-pQTLs associated with the target protein annotated and effect estimates were aligned to the protein increasing allele, i.e. bars with a positive –log10(p-values) indicate positive associations with a trait from the database and *vice versa*. Clinical traits are grouped by higher-level categories and coloured accordingly. GIT = gastrointestinal tract, Misc = Miscellaneous, No coloc. pos. = colocalisation for secondary signals was not possible

We observed general consistency among phenotypic traits colocalising with *cis*-pQTLs, i.e. traits were closely related and effect estimates were consistent with phenotypic presentations (**Supplemental Tab. S7 and Fig. 6**). For instance, rs165656, a lead *cis*-pQTL increasing catechol o-methyltransferase plasma abundances, is a regional lead variant for BMI^32^ and specifically colocalised with adiposity related traits, i.e. inversely associated with overall measures of body size such as BMI, weight, and fat-free mass. In general, phenotypic characterization of potential genetic instruments to simulate targeting abundances or activities of proteins can help to distinguish those with narrow and well-defined or target-specific from those with undesirable or broad phenotypic effects. Notable exceptions included the IL-6 receptor variant rs2228145, for which the protein increasing C allele was inversely associated with the risk of coronary heart disease and rheumatoid arthritis but positively with the risk for allergic disease, such as asthma.

### A variant at the ABO locus links susceptibility of respiratory failure in COVID-19 to protein targets

A recent GWAS identified two independent genomic loci to be associated with an increased risk of respiratory failure in COVID-19 patients^10^. We observed six proteins to be associated positively with the lead signal (rs657152) at the *ABO* locus (coagulation factor VIII, sulfhydryl oxidase 2 (QSOX2), von Willebrand factor, SVEP1, and heme oxygenase 1) and one inverse association (interleukin-6 receptor subunit beta), but did not observe significantly associated proteins with the lead variant (rs11385942) at 3p21.31. We identified a cluster of ten aptamers (targeting SVEP1, coagulation factor VIII, ferritin, heme oxygenase 1, van Willebrand factor, plasminogen, PLOD2, and CD14) sharing a genetic signal (regional probability: 0.88; rs941137; **Supplemental Fig. S4**), which was in high LD (r^2^=0.85) with the lead *ABO* signal associated with a higher risk for respiratory failure among COVID-19 patients.

### Webserver

To facilitate in-depth exploration of candidate proteins, i.e. those with at least one *cis*-pQTL, we created an online resource (https://omicscience.org/apps/covidpgwas/). The webserver provides an intuitive representation of genetic findings, including the opportunity of customized look-ups and downloads of the summary statistics for specific genomic regions and protein targets of interest. We further provide detailed information for each protein target, including links to relevant databases, such as UniProt or Reactome, information on currently available drugs or those in development as well as characterization of associated SNPs. The webserver further enables the query of SNPs across proteins to assess specificity and to find co-associated protein targets.

## DISCUSSION

We present the largest and most systematic genetic investigation of host proteins reported to interact with SARS-CoV-2 proteins, be related to virus entry, host hyperimmune or procoagulant responses, or be associated with the severity of COVID-19. The integration of large-scale genomic and aptamer-based plasma proteomic data from 10,708 individuals improves our understanding of the genetic architecture of 97 of 179 investigated host proteins by identifying 220 *cis-*acting variants that explain up to 70% of the variance in these proteins, including 45 with no previously known pQTL and 38 encoding current drug targets. Our findings, shared in an interactive webserver (https://omicscience.org/apps/covidpgwas/), enable rapid ‘in silico’ follow-up of these variants and assessment of their causal relevance as molecular targets for new or repurposed drugs in human genetic studies of SARS-CoV-2 and COVID-19, such as the COVID-19 Host Genetics Initiative (https://www.covid19hg.org/).

The contribution of identified genetic variants outweighed the variance explained by most of the tested host factors for the majority of protein targets. Protein expression in plasma was also frequently associated with expression of protein encoding genes in relevant tissues. We demonstrate that a large number of genetic variants acting in *trans* are non-specific and show evidence of substantial horizontal pleiotropy. Findings for these variants should be treated with caution in follow-up studies focused on protein-specific genetic effects.

The successful identification of druggable targets for COVID-19 provides an insight both on potential therapies but also on medications that might worsen outlook, depending on the direction of the genetic effect, and whether any associated compound inhibits or activates the target. We also found genetic evidence that selected protein targets, such as for MARK3 and monocyte count, have potential for adverse effects on other health outcomes, but note that this was not a general characteristic of all tested ‘druggable’ targets. Further, in-depth characterization of the targets identified will be required as a first step in gauging the likely success of any new or repurposed drugs identified via this analysis^33^.

We exemplify the value of the data resource generated by being the first that links a genomic risk variant for poor prognosis among COVID-19 patients, i.e. respiratory failure, at the *ABO* locus^10^ to proteins related to the maladaptive response of the host, namely hypercoagulation, as well as two putative viral interaction partners (heme oxygenase 1 and PLOD2). The risk increasing A allele of rs657152 was consistently associated with higher plasma levels of coagulation factor VIII and von Willebrand factor. Anticoagulation is associated with a better outcome in patients with severe COVID-19^34^, and randomised controlled trails are underway to properly evaluate the benefit or harms of anticoagulant therapies.

Affinity-based proteomics techniques rely on conserved binding epitopes. Changes in the 3D-conformational structure of target proteins introduced by protein altering variants (PAVs) might change the binding affinity to the target, and hence measurements, without affecting biological activity of the protein. We identified 52 *cis*-pQTLs which were in LD (r^2^>0.1) with a PAV. However, 27 of those *cis*-pQTLs or a proxy in high LD (r^2^>0.8) have been previously identified as genome-wide significant signals for at least one trait in the GWAS catalogue (excluding any entries of platforms used in the present study) and might therefore carry biologically meaningful information.

This study is the largest genetic discovery of protein targets highly relevant to the current COVID-19 pandemic and was designed to provide a rapid open access platform to help prioritise drug discovery and repurposing efforts. However, important limitations apply. Firstly, protein abundances have been measured in plasma, which may differ from the intracellular role of proteins, and include purposefully secreted as well as leaked proteins. Secondly, while aptamer-based techniques provide the broadest coverage of the plasma proteome, specificity can be compromised for specific protein targets and evidence using complementary techniques such as Olink or mass spectrometry efforts is useful for validation of signals. Thirdly, in-depth phenotypic characterization of the high-priority *cis*-pQTLs requires appropriate formal and statistical follow-up, such as colocalisation, where the genomic architecture permits existing approaches not yet optimised for multiple secondary signals and outcomes, and *cis*-GRS evaluation in independent and adequately powered studies for the trait of interest.

## Materials and Methods

### Study participants

The Fenland study is a population-based cohort of 12,435 participants born between 1950 and 1975 who underwent detailed phenotyping at the baseline visit from 2005-2015. Participants were recruited from general practice surgeries in the Cambridgeshire region in the UK. Exclusion criteria were: clinically diagnosed diabetes mellitus, inability to walk unaided, terminal illness, clinically diagnosed psychotic disorder, pregnancy or lactation. The study was approved by the Cambridge Local Research Ethics Committee (ref. 04/Q0108/19) and all participants provided written informed consent. Population characteristics and proteomic measures have previously been described in detail^35^.

### Mapping of protein targets across platforms

We mapped each candidate protein to its UniProt-ID (https://www.uniprot.org/) and used those to select mapping aptamers and Olink measures based on annotation files provided by the vendors.

### Proteomic profiling

Proteomic profiling of fasted EDTA plasma samples from 12,084 Fenland Study participants collected at baseline was performed by SomaLogic Inc. (Boulder, US) using an aptamer-based technology (SOMAscan proteomic assay). Relative protein abundances of 4,775 human protein targets were evaluated by 4,979 aptamers (SomaLogic V4), as previously described^35^. To account for variation in hybridization within runs, hybridization control probes are used to generate a hybridization scale factor for each sample. To control for total signal differences between samples due to variation in overall protein concentration or technical factors such as reagent concentration, pipetting or assay timing, a ratio between each aptamer’s measured value and a reference value is computed, and the median of these ratios is computed for each of the three dilution sets (40%, 1% and 0.005%) and applied to each dilution set. Samples were removed if they were deemed by SomaLogic to have failed or did not meet our acceptance criteria of 0.25-4 for all scaling factors. In addition to passing SomaLogic QC, only human protein targets were taken forward for subsequent analysis (4,979 out of the 5284 aptamers). Aptamers’ target annotation and mapping to UniProt accession numbers as well as Entrez gene identifiers were provided by SomaLogic.

Plasma samples for a subset of 500 Fenland participants were additionally measured using 12 Olink 92-protein panels using proximity extension assays^36^. Of the 1104 Olink proteins, 1069 were unique (n=35 on >1 panel, average correlation coefficient 0.90). We imputed values below the detection limit of the assay using raw fluorescence values. Protein levels were normalized (‘NPX’) and subsequently log2-transformed for statistical analysis. A total of 15 samples were excluded based on quality thresholds recommended by Olink, leaving 485 samples for analysis.

### Genotyping and imputation

Fenland participants were genotyped using three genotyping arrays: the Affymetrix UK Biobank Axiom array (OMICs, N=8994), Illumina Infinium Core Exome 24v1 (Core-Exome, N=1060) and Affymetrix SNP5.0 (GWAS, N=1402). Samples were excluded for the following reasons: 1) failed channel contrast (DishQC <0.82); 2) low call rate (<95%); 3) gender mismatch between reported and genetic sex; 4) heterozygosity outlier; 5) unusually high number of singleton genotypes or 6) impossible identity-by-descent values. Single nucleotide polymorphisms (SNPs) were removed if: 1) call rate < 95%; 2) clusters failed Affymetrix SNPolisher standard tests and thresholds; 3) MAF was significantly affected by plate; 4) SNP was a duplicate based on chromosome, position and alleles (selecting the best probeset according to Affymetrix SNPolisher); 5) Hardy-Weinberg equilibrium p<10^−6^; 6) did not match the reference or 7) MAF=0.

Autosomes for the OMICS and GWAS subsets were imputed to the HRC (r1) panel using IMPUTE4^37^, and the Core-Exome subset and the X-chromosome (for all subsets) were imputed to HRC.r1.1 using the Sanger imputation server (https://imputation.sanger.ac.uk/)^38^. All three arrays subsets were also imputed to the UK10K+1000Gphase3^39^ panel using the Sanger imputation server in order to obtain additional variants that do not exist in the HRC reference panel. Variants with MAF < 0.001, imputation quality (info) < 0.4 or Hardy Weinberg Equilibrium p < 10^−7^ in any of the genotyping subsets were excluded from further analyses.

### GWAS and meta-analysis

After excluding ancestry outliers and related individuals, 10,708 Fenland participants had both phenotypes and genetic data for the GWAS (OMICS=8,350, Core-Exome=1,026, GWAS=1,332). Within each genotyping subset, aptamer abundances were transformed to follow a normal distribution using the rank-based inverse normal transformation. Transformed aptamer abundances were then adjusted for age, sex, sample collection site and 10 principal components and the residuals used as input for the genetic association analyses. Test site was omitted for protein abundances measured by Olink as those were all selected from the same test site. Genome-wide association was performed under an additive model using BGENIE (v1.3)^37^. Results for the three genotyping arrays were combined in a fixed-effects meta-analysis in METAL^40^. Following the meta-analysis, 17,652,797 genetic variants also present in the largest subset of the Fenland data (Fenland-OMICS) were taken forward for further analysis.

### Definition of genomic regions (including cis/trans)

For each aptamer, we used a genome-wide significance threshold of 5×10^−8^ and defined non-overlapping regions by merging overlapping or adjoining 1Mb intervals around all genome-wide significant variants (500kb either side), treating the extended MHC region (chr6:25.5–34.0Mb) as one region. For each region we defined a regional sentinel variant as the most significant variant in the region. We defined genomic regions shared across aptamers if regional sentinels of overlapping regions were in strong LD (r^2^>0.8).

### Conditional analysis

We performed conditional analysis as implemented in the GCTA software using the *slct* option for each genomic region - aptamer pair identified. We used a collinear cut-off of 0.1 and a p-value below 5×10^−8^ to identify secondary signals in a given region. As a quality control step, we fitted a final model including all identified variants for a given genomic region using individual level data in the largest available data set (‘Fenland-OMICs’) and discarded all variants no longer meeting genome-wide significance.

We performed a forward stepwise selection procedure to identify secondary signals at each locus on the X-chromosome using SNPTEST v.2.5.2 to compute conditional GWAS based on individual level data in the largest subset. Briefly, we defined conditionally independent signals as those emerging after conditioning on all previously selected signals in the locus until no signal was genome-wide significant.

### Explained variance

To compute the explained variance for plasma abundancies of protein targets we fitted linear regression models with residual protein abundancies (see GWAS section) as outcome and 1) only the lead *cis*-pQTL, 2) all *cis*-pQTLs, or 3) all identified pQTLs as exposure. We report the R^2^ from those models as explained variance.

### Annotation of pQTLs

For each identified pQTL we first obtained all SNPs in at least moderate LD (r^2^>0.1) and queried comprehensive annotations using the variant effect predictor software^41^ (version 98.3) using the *pick* option. For each *cis*-pQTL we checked whether either the variant itself or a proxy in the encoding gene (r^2^>0.1) is predicted to induce a change in the amino acid sequence of the associated protein, so-called protein altering variants (PAVs).

### Mapping of cis-pQTLs to drug targets

To annotate druggable targets we merged the list of proteins targeted by the SomaScan V4 platform with the list of druggable genes from Finan at al.^13^ based on common gene entries. We further added protein – drug combinations as recommended by Gordon et al.^3^.

### Identification of relevant GWAS traits

To enable linkage to reported GWAS-variants we downloaded all SNPs reported in the GWAS catalog (19/12/2019, https://www.ebi.ac.uk/gwas/) and pruned the list of variant-outcome associations manually to omit previous protein-wide GWAS. For each SNP identified in the present study (N=671) we tested whether the variant or a proxy in LD (r^2^>0.8) has been reported to be associated with other outcomes previously.

### Definition of novel pQTLs

To test whether any of the identified regional sentinel pQTLs has been reported previously, we obtained a list of published pQTLs^15,24,26,42,43^ and defined novel pQTLs as those not in LD (r^2^<0.1) with any previously identified variant. We note that this approach is rather conservative, since it only asks whether or not any of the reported SNPs has ever been reported to be associated with any protein measured with multiplex methods.

### Assessment of pleiotropy

To evaluate possible protein-specific pleiotropy of pQTLs we computed association statistics for each of the 671 unique SNPs across 4,979 aptamers (N=4,775 unique protein targets) with the same adjustment set as in the GWAS. This resulted in a protein profile for each variant defined as all aptamers significantly associated (p<5×10^−8^). For all aptamers we retrieved all GO-terms referring to biological processes from the UniProt database using all possible UniProt-IDs as a query. GO-term annotation within the UniProt database has the advantage of being manually curated while aiming to omit unspecific parent terms. We tested for each pQTL if the associated aptamers fall into one of the following criteria: 1) solely associated with a specific protein, 2) all associated aptamers belong to a single GO-term, 3) the majority (>50%) of associated aptamers but at least two belong to a single GO-term, and 4) no single GO-term covers more than 50% of the associated aptamers. We refer to category 1 as protein-specific association, categories 2 and 3 as vertical pleiotropy, and category 4 as horizontal pleiotropy.

### Heritability estimates and genetic correlation

We used genome-wide genotype data from 8,350 Fenland participants (Fenland-OMICs) to determine SNP-based heritability and genetic correlation estimates among the 102 protein targets with at least one *cis*-pQTLs and excluding proteins encoded in the X-chromosome. We generated a genetic relationship matrix (GRM) using GCTA v.1.90^44^ from all variants with MAF > 1% to calculate SNP-based heritability as implemented by biMM^45^. Genetic correlations were computed between all 4273 possible pairs among 93 protein targets with heritability estimates larger than 1.5 times its standard error, using the generated GRM by a bivariate linear mixed model as implemented by biMM. We further conducted two sensitivity analyses to evaluate whether the estimated genetic correlation could be largely attributable to the top *cis*-pQTL or to shared pleiotropic *trans* regions. To evaluate contribution of the top *cis* variant, each protein target was regressed against its sentinel *cis* variant in addition to age, sex, sample collection site, 10 principal components and the residuals were used as phenotypes to compute heritability and genetic correlation estimates. To assess the contribution of 29 pleiotropic *trans* regions, we excluded 2Mb genomic regions around pleiotropic *trans*-pQTLs (associated with >20 aptamers) from the GRM to compute heritability and genetic correlation estimates. Genetic correlations could not be computed for pairs involving IL1RL1 in the main analysis and were therefore excluded. However, upon regressing out the sentinel *cis*-variant, genetic correlations with this protein could be computed probably due to its large contribution to heritability.

### Variance decomposition

We used linear mixed models as implemented in the R package *variancePartition* to decompose inverse rank-normal transformed plasma abundances of 106 aptamers with at least one *cis*-pQTL. To this end, we computed weighted genetic scores for each aptamer separating SNPs acting in *cis* (*cis*-GRS) and *trans* (*trans*-GRS). In addition to the GRS we used participants’ age, sex, body mass index, waist-to-hip ratio, systolic and diastolic blood pressure, reported alcohol intake, smoking consumption and fasting plasma levels of glucose, insulin, high-density lipoprotein cholesterol, low-density lipoprotein cholesterol, alanine aminotransaminase as well as a creatinine-based estimated glomerular filtration rate as explanatory factors. We implemented this analysis in the Fenland-OMICs data set leaving 8,004 participants without any missing values in the factors considered.

### Genetic risk scores associations

We computed weighted GRS for metabolic (Insulin resistance^46^, type 2 diabetes^47^ and BMI^48^), respiratory (forced expiratory volume, forced vital capacity^19^ and asthma^49^) and cardiovascular traits (eGFR^50^, systolic blood pressure^51^, diastolic blood pressure^51^ and coronary artery disease^30^) for Fenland-OMICs participants (N = 8,350) to evaluate their association with plasma protein abundances. GRSs were computed from previously reported genome-wide significant variants and weighted by their reported beta coefficients for continuous outcomes or log(OR) for binary outcomes. Variants not available among Fenland genotypes, strand ambiguous or with low imputation quality (INFO < 0.6) were excluded from the GRSs. Associations between each scaled GRS and log10 transformed and scaled protein levels were computed by linear regressions adjusted by age, sex, 10 genetic principal components and sample collection site.

We implemented this analysis for the 186 proteins with at least one associated cis or trans-pQTL. Associations with p-values < 0.05/186 were deemed significant according to Bonferroni correction for multiple comparisons.

### Incorporation of GTEx v8 data

We leveraged gene expression data in five human tissues (lung, whole blood, heart - left ventricle, heart - atrial appendage, and liver), of relevance to COVID-19 and its potential adverse effects and complications, from the Genotype-Tissue Expression (GTEx) project^21,22^. For the 102 Somamers with at least one *cis*-pQTL located on the autosomes and available gene expression models trained in GTEx v8^52^, we performed summary-statistics based PrediXcan^23^ analysis to identify tissue-dependent genetically determined gene expression traits that significantly predict plasma protein levels. We used the standardized effect size (*z*-score) to investigate the tissue specificity or the consistency of the association across the tissues between the genetic component of the expression of the encoding gene and the corresponding protein. We performed DAVID functional enrichment analyses on all the genes significantly associated (Bonferroni-adjusted p<0.05) with plasma levels of the proteins to identify biological processes (Benjamini-Hochberg adjusted p<0.05) that may explain the associations found beyond the protein encoding genes.

### Cross-platform comparison

We selected 24 *cis*- and 101 *trans*-pQTLs mapping to 33 protein targets overlapping with Olink from the SomaScan-based discovery and obtained summary statistics from in-house genome-wide association studies (GWAS) based on corresponding Olink measures. To enable a more systematic reciprocal comparison, we further compared 13 pQTLs (for 11 proteins) only apparent in an in-house Olink-based pGWAS (p<4.5×10^−11^) effort and obtained GWAS-summary statistics from corresponding aptamer measurements. We pruned the list for variants in high LD (r^2^>0.8) and discarded SNPs not passing QC for both efforts (n=6).

### Phenome-wide scan among UK Biobank and look-up

We obtained all ICD-10 codes-related genome-wide summary statistics from the most recent release of the Neale lab (http://www.nealelab.is/uk-biobank) with at least 100 cases resulting in 633 distinct ICD-10 codes. Among the 220 *cis*-pQTLs identified in the present study, 215 were included in the UK Biobank summary statistics (3 aptamers had to be excluded due to unavailable lead *cis*-pQTLs or proxies in LD). We next aligned effect estimates between *cis*-pQTLs and UK Biobank statistics and used the *grs*.*summary()* function from the ‘gtx’ R package to compute the effect of a weighted *cis*-GRS for an aptamer across all 633 ICD-codes. We applied a global testing correction across all cis-GRS – ICD-10 code combinations using the Benjamini-Hochberg procedure and declared a false discovery rate of 10% as a significance threshold.

We queried all 220 *cis*-pQTLs for genome-wide association results using the *phewas()* function of the R package ‘ieugwasr’ linked to the IEU GWAS database. We selected all variants in strong LD (r^2^>0.8) with any of the *cis*-pQTLs to incorporate information on proxies. We restricted the search in the ieugwar tool to the batches “ebi-a“, “ieu-a“, and “ukb-b” to minimize redundant phenotypes.

### Colocalisation analysis

We used statistical colocalisation^53^ to test for a shared genetic signal between a protein target and a phenotype with evidence of a significant effect of the *cis*-pQTL (see above). We obtained posterior probabilities (PP) of: H0 – no signal; H1 – signal unique to the protein target; H2 – signal unique to the trait; H3 – two distinct causal variants in the same locus and H4 – presence of a shared causal variant between a protein target and a given trait. PPs above 75% were considered highly likely. In case the *cis*-pQTL was a secondary signal we computed conditional association statistics using the *cond* option from GCTA-cojo to align with the identification of secondary signals. We conditioned on all other secondary signals in the locus. We note that conditioning on all other secondary variants in the locus failed to produce the desired conditional association statistics in a few cases probably due to moderate LD (r^2^>0.1) between selected secondary variants and other putative secondary variants.

### Multi-trait colocalization at the ABO locus

We used hypothesis prioritisation in multi-trait colocalization (HyPrColoc)^54^ at the *ABO* locus (±200kb) 1) to identify protein targets sharing a common causal variant over and above what could be identified in the meta-analysis to increase statistical power, and 2) to identify possible multiple causal variants with distinct associated protein clusters. Briefly, HyPrColoc aims to test the global hypothesis that multiple traits share a common genetic signal at a genomic location and further uses a clustering algorithm to partition possible clusters of traits with distinct causal variants within the same genomic region. HyPrColoc provides for each cluster three different types of output: 1) a posterior probability (PP) that all traits in the cluster share a common genetic signal, 2) a regional association probability, i.e. that all the metabolites share an association with one or more variants in the region, and 3) the proportion of the PP explained by the candidate variant. We considered a highly likely alignment of a genetic signal across various traits if the regional association probability > 80%. This criterion takes to some extend into account that metabolites may share multiple causal variants at the same locus and provides some robustness against violation of the single causal variant assumption. We note that several protein targets had multiple independent signals at the ABO locus (**Supplementary Tab. S4**). We further filtered protein targets with no evidence of a likely genetic signal (p>10^−5^) in the region before performing HyPrColoc, which improved clustering across traits due to minimizing noise.

## ACKNOWLEDGMENTS AND FUNDING

The Fenland Study (10.22025/2017.10.101.00001) is funded by the Medical Research Council (MC_UU_12015/1). We are grateful to all the volunteers and to the General Practitioners and practice staff for assistance with recruitment. We thank the Fenland Study Investigators, Fenland Study Co-ordination team and the Epidemiology Field, Data and Laboratory teams. We further acknowledge support for genomics from the Medical Research Council (MC_PC_13046). Proteomic measurements were supported and governed by a collaboration agreement between the University of Cambridge and Somalogic. JCZ and VPWA are supported by a 4-year Wellcome Trust PhD Studentship and the Cambridge Trust, CL, EW, and NJW are funded by the Medical Research Council (MC_UU_12015/1). NJW is an NIHR Senior Investigator. GK is supported by grants from the National Institute on Aging (NIA): R01 AG057452, RF1 AG058942, RF1 AG059093, U01 AG061359, and U19 AG063744. MR acknowledges funding from the Francis Crick Institute, which receives its core funding from Cancer Research UK (FC001134), the UK Medical Research Council (FC001134), and the Wellcome Trust (FC001134). ERG is supported by the National Institutes of Health Genomic Innovator Award (<u>R35 HG010718</u>). JR is supported by the German Federal Ministry of Education and Research (BMBF) within the framework of the e:Med research and funding concept (grant no. 01ZX1912D).

## AUTHOR CONTRIBUTIONS

MP, ADH, and CL designed the analysis and drafted the manuscript. MP, EW, JCSZ, VPWA, and JL analysed the data. NK and EO performed quality control of proteomic measurements. JR and GK designed and implemented the webserver. RO and SW advised proteome measurements and assisted in quality control. EG did the gene expression analysis and interpretation of results. JPC and MR provided critical review and intellectual contribution to the discussion of results. NJW is PI of the Fenland cohort. All authors contributed to the interpretation of results and critically reviewed the manuscript.

## DATA AVAILABILITY

All genome-wide summary statistics are made available through an interactive webserver (https://omicscience.org/apps/covidpgwas/).

## CODE AVAILABILITY

Each use of software programs has been clearly indicated and information on the options that were used is provided in the Methods section. Source code to call programs is available upon request.

