## Supplemental Material for "Genetic architecture of host proteins interacting with SARS-CoV-2"

### FIGURES

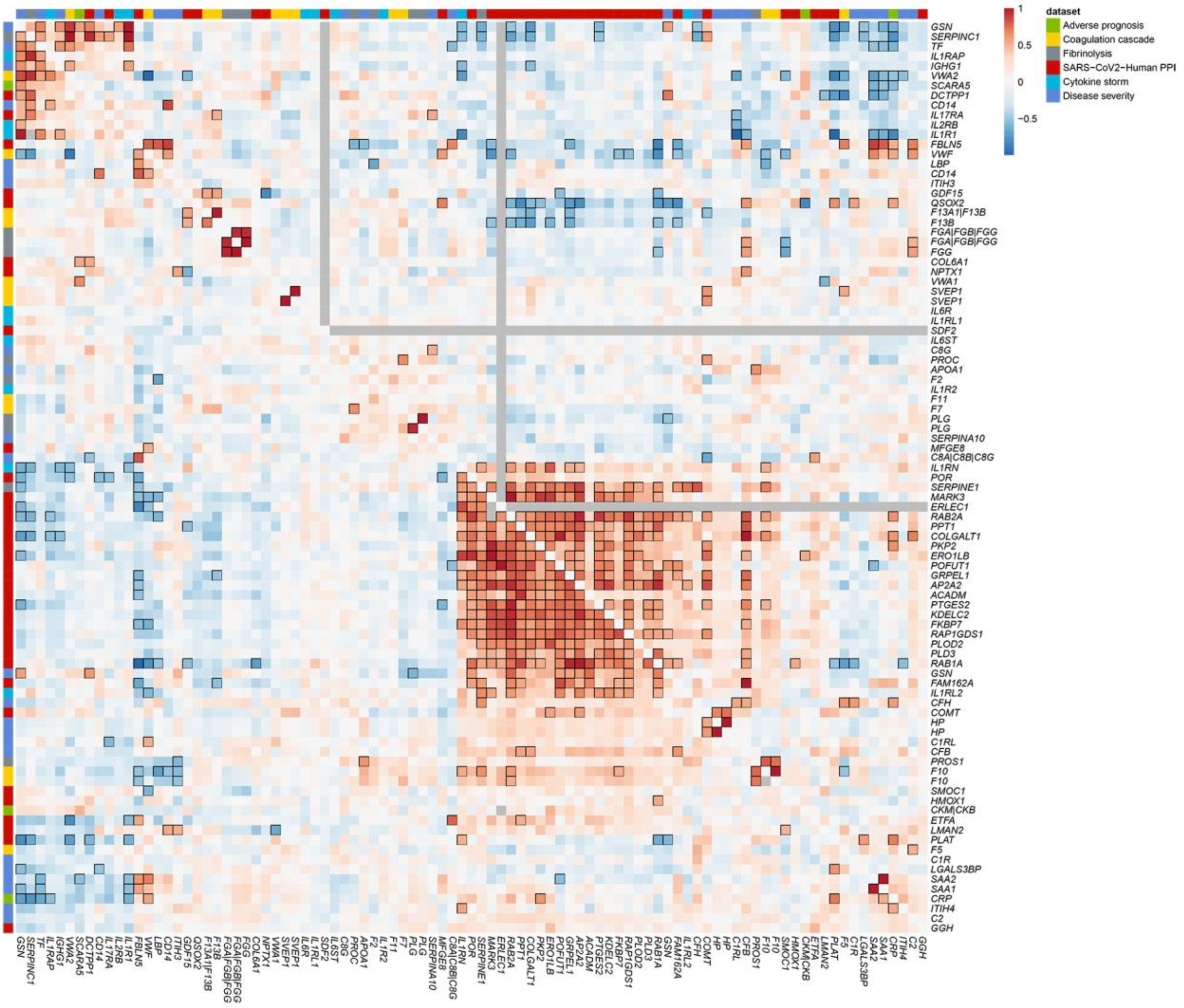


**Figure S1** Genetic correlation of 93 Somamers regressing out the top sentinel cis-pQTL (lower triangle) and excluding pleiotropic trans genomic regions (upper triangle). SOMAmers were clustered based on absolute genetic correlations and protein encoding genes were used as labels. The column on the far left indicates relevance to SARS-CoV2 infection. Strong correlations (|r|>0.5) are indicated by black frames. Values for proteins with low heritability estimates (heritability < 1.5 x SE(heritability)) in these sensitivity analyses are shown in grey.

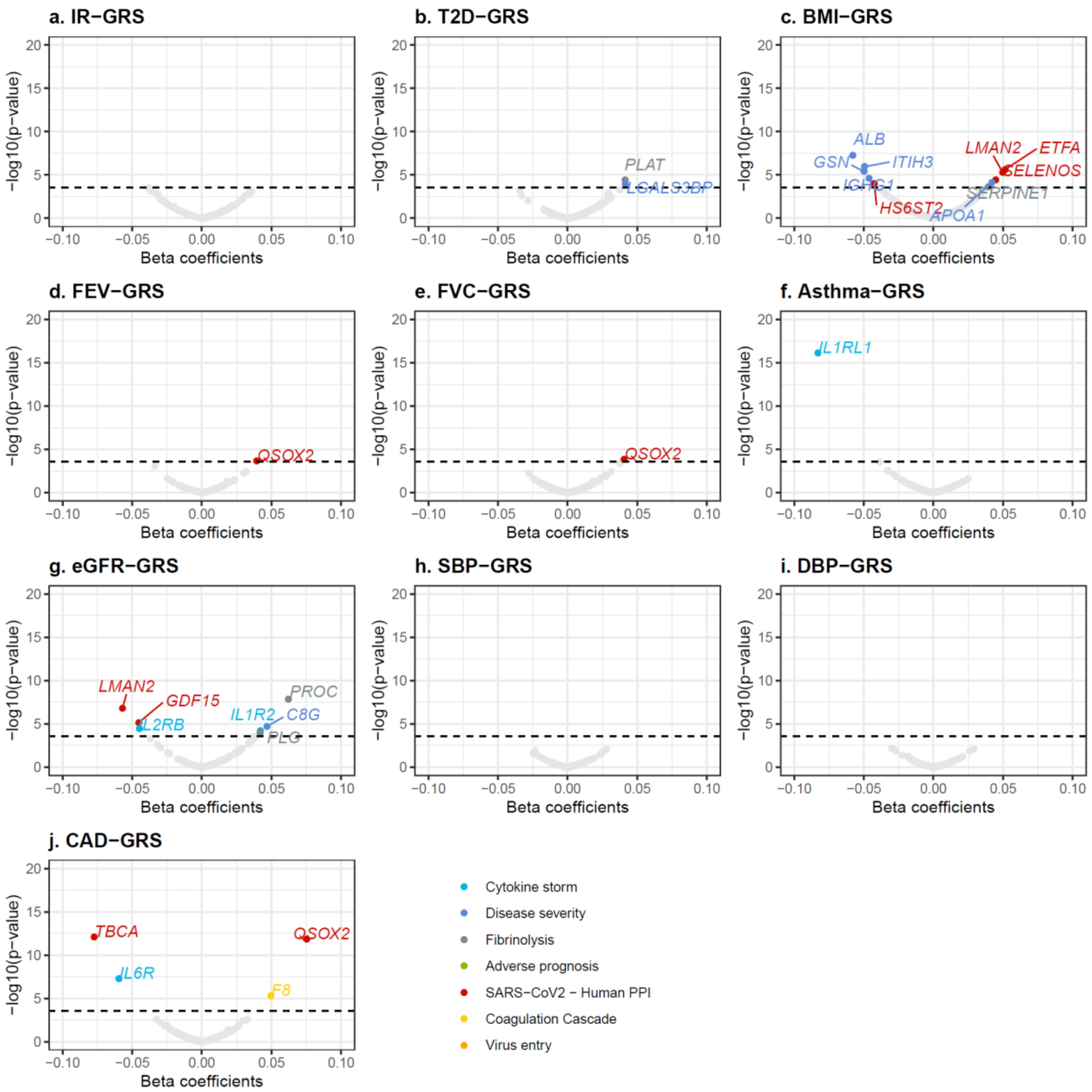


**Figure S2** Differences in plasma abundances of 185 Somamers targeting candidate proteins by genetic susceptibility to diseases and risk phenotypes. Each panel shows the association of genetic risk scores for metabolic (T2D = type 2 diabetes, IR = insulin resistance, BMI = body mass index), respiratory (FEV = forced expired volume, FVC = forced vital capacity, Asthma) and cardiovascular (eGFR = estimated glomerular filtration rate, SBP = systolic blood pressure, DBP = diastolic blood pressure, CAD = coronary artery disease) phenotypes with protein levels. Significant associations (p-value < 0.05/186 for multiple comparisons) are coloured according to COVID-19 associated processes.

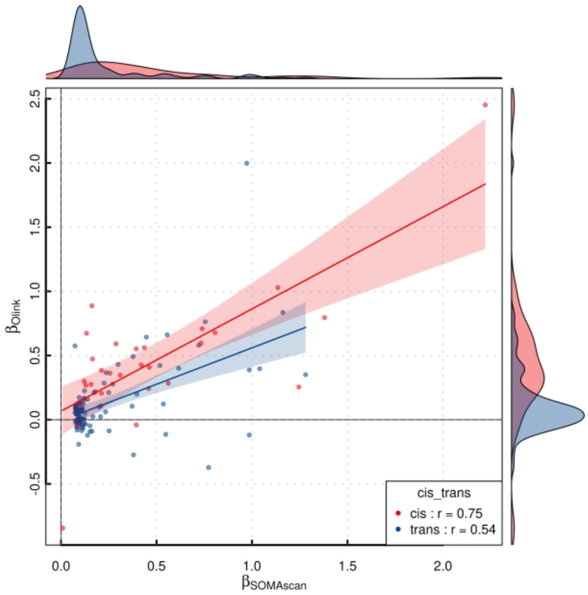


**Figure S3** Comparison of effect estimates for protein quantitative loci (pQTL) discovered using either the SOMAscan or Olink platforms for 32 proteins targeted by both platforms. Colours indicate the proximity to the protein encoding gene (red - *cis*, blue - *trans*). Correlation coefficients (r) are given stratified by cis/trans status. Effect estimates have been aligned to be positive for all aptamer associations to enable unbiased assessment of the correlation coefficient.

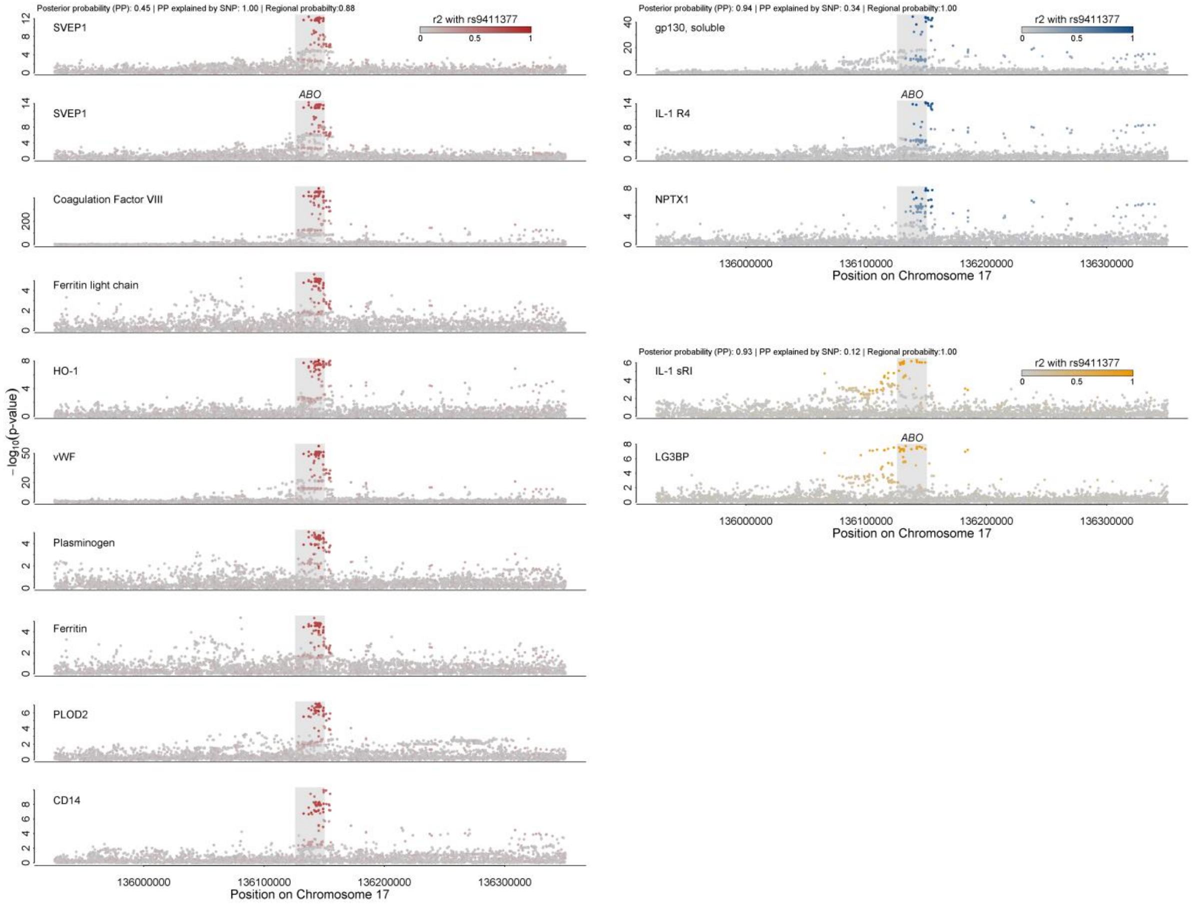


**Figure S4** Regional association plots centred on the *ABO* locus. Multi-trait colocalization identified three cluster of protein associations distributed across three distinct likely causal variants. P-values are coloured based on linkage disequilibrium with the candidate variant prioritized by the colocalization algorithm (red, blue, and orange). Probabilities are given on top of each cluster.
